# XWRAP^®^ Human Amniotic Membrane in Combination with an Amniotic-Fluid-Derived Biologic Promotes Collagen Fibril Regeneration in a Preclinical Rabbit Model of Medial Collateral Ligament Injury

**DOI:** 10.64898/2026.09.18.752432

**Authors:** Joydeep Basu

## Abstract

Current treatment of orthopedic ligament injury relies principally on non-regenerative materials and surgical repair rather than strategies that promote intrinsic tissue regeneration. This study evaluated whether biologic augmentation with XWRAP^®^, a human amniotic membrane allograft, combined with a cryopreserved amniotic-fluid-derived biologic containing micronized human amnion membrane, promotes regeneration of ligament architecture in a rabbit medial collateral ligament (MCL) gap-defect model. Standardized 4-mm mid-substance MCL gap defects were created in New Zealand White rabbits (n=3) and either treated with the amniotic membrane/amniotic-fluid biologic combination or left untreated, with intact contralateral or adjacent MCL serving as native controls; ligaments were evaluated by transmission electron microscopy at 2 and 6 weeks post-procedure, with collagen fibril diameter quantified from calibrated high-magnification micrographs. Treated defects developed significantly larger mean collagen fibril diameters than untreated defects at both time points (75 vs. 50 nm at 2 weeks and 88 vs. 62 nm at 6 weeks; p<0.0001 for each comparison), reaching approximately 90% of native, uninjured ligament fibril diameter (98 nm) by 6 weeks. Treated defects also developed a broader, bimodal fibril-diameter distribution resembling native ligament architecture, whereas untreated defects remained characterized by a narrow, unimodal population of small-diameter fibrils typical of scar-associated repair. These findings indicate that biologic augmentation with XWRAP^®^ combined with an amniotic-fluid-derived biologic promotes a regenerative, rather than purely reparative, healing response following ligament injury, progressively restoring collagen fibril architecture toward that of native tissue in this preclinical model.

## Introduction

Ligament injuries represent a substantial and persistent burden in orthopedic practice, and the medial collateral ligament (MCL) of the knee is among the most frequently injured stabilizing structures of the musculoskeletal system [4]. Unlike many other connective tissues, ligaments possess a limited intrinsic capacity for true regeneration; when disrupted, they typically heal through a fibrovascular scarring process that yields mechanically and structurally inferior tissue rather than native ligament architecture [4]. This distinction between reparative (scar) healing and true regeneration is most clearly evident at the level of the collagen fibril: normal, mature ligament is characterized by a bimodal distribution of large- and small-diameter collagen fibrils, whereas scar tissue formed during conventional healing is dominated by a unimodal population of small-diameter fibrils [1–3] that correlates with reduced collagen crosslink density and diminished mechanical strength [5]. These fibril-level differences, first systematically characterized in the rabbit MCL model [1–3], remain a well-established structural benchmark for distinguishing regenerative from reparative healing outcomes and are the structural endpoint examined in the present study [6].

Current standard of care for ligament and other orthopedic soft-tissue trauma relies principally on non-regenerative materials and surgical techniques that stabilize or reconstruct the injured tissue without addressing the underlying biological deficit that drives scar-mediated repair. In recent years, biologic augmentation — the use of cell-based or acellular biological materials to actively promote a regenerative, rather than purely reparative, healing response — has emerged as an active area of investigation across multiple ligament injury applications, including the anterior cruciate ligament, reflecting broader interest in shifting the therapeutic paradigm from repair toward regeneration [7].

Human amniotic membrane and amniotic fluid-derived products are of particular interest as biologic augmentation materials given their naturally high content of structural extracellular matrix proteins together with bioactive cytokines and growth factors that support tissue remodeling [8]. Amniotic tissue-derived products have already seen expanding clinical use across a range of orthopedic applications, including tendon and ligament injury, cartilage defects, and osteoarthritis, with early clinical and preclinical evidence supporting their safety and regenerative potential [9]. Applied Biologics’ biomaterial compound, which embeds multiple bioactive proteins, cytokines, and growth factors within an extracellular matrix scaffold derived from human amnion, was developed to leverage these properties directly at the site of ligament injury.

In this study, we evaluated a biologic augmentation strategy combining XWRAP^®^, a human amniotic membrane allograft, with an amniotic-fluid-derived biologic containing micronized human amniotic membrane, in a preclinical rabbit model of MCL gap-defect injury. Building on the established use of collagen fibril diameter distribution as a structural marker of regenerative versus reparative ligament healing [1–3,5], we assessed whether this combination biomaterial shifts the healing trajectory of the injured MCL toward a native-like, regenerative ultrastructural phenotype, rather than the small-diameter, scar-like fibril population characteristic of untreated healing.

## Materials and Methods

### 1) Preclinical animal model

Species: *Oryctolagus cuniculus*

Strain: New Zealand White, 12-month-old

### 2) Study design

#### Pre-pilot. *n*=1. Rabbit ID #603

A 4 mm mid-substance segment of the left and right hind-limb MCL was surgically removed. The surgical site was marked by sewing four 6-0 monofilament nylon sutures at the remnant MCL corners. For the right MCL (experimental), a small piece of XWRAP^®^ was cut to size and sutured over the injury site and 1-2 ml cryopreserved human amniotic fluid with micronized human amnion membrane biomaterial compound was injected under the membrane. The left hind-limb MCL serves as an untreated control. Injury sites were surgically closed and animal was allowed unrestricted cage activity post-operation. Although the overall study duration was 6 weeks, this animal was humanely sacrificed at 2 weeks post-surgery and the left/right MCL removed. Normal (native) control MCL for this animal was sourced from MCL adjacent to the injury site. The regenerated/healed area was identified by the region between the sutures; this area was evaluated by transmission electron microscopy for density and distribution of regenerated collagen fibrils.

#### Pilot. n=2

One animal is experimental (**Rabbit ID #608**), one animal is control (**Rabbit ID #740**). For both animals, the right MCL was resected as above. In the experimental animal, treatment was provided as described above, for the control animal no treatment was provided. Study duration is 6 weeks post-surgery. In both animals, the left hind-limb MCL becomes an intact, uninjured control comparator for TEM evaluation.

Each rabbit had a standardized surgical injury created on the medial collateral ligament (MCL) of the right hind-limb (or both right and left hind-limbs for the pre-pilot animal). This consists of removal of a mid-substance segment of the MCL, leaving a 4.0 mm +/-0.5 mm gap in the ligament with the knee at 70 degrees +/-10 degrees of flexion. In order to mark the area of the defect clearly, four 6-0 monofilament nylon sutures were sewn into the corners of the ligament. For control animals, the wound site was closed without further intervention. For experimental animals, a small piece of human amniotic membrane XWRAP^®^ is sutured over the injury site. 1-2 ml of cryopreserved human amniotic fluid with micronized human amnion membrane biomaterial compound is injected under the membrane. The wound site is closed and the animal transitioned to standard post-op recovery and care.

### 3) Preparation of tissue for electron microscopy

Tissue was prepared as described in [1-3]. Briefly, the left and right medial collateral ligaments (MCL) from rabbit ID No. 603, 608 and 740 were carefully dissected and fixed in 10% formalin for at least 48 hours prior to submission for transmission electron microscopy (TEM). Using a razor blade, longitudinal strips (3 mm x 5 mm) were collected from the center mid-substance from each MCL and further subdivided into 1 mm x 1 mm strips and placed in Karnovsky’s fixative for subsequent cross-sectional TEM evaluation. Using a Zeiss EM 910 TEM microscope, multiple TEM digital micrographs of fibril cross-sections were captured at 2.5kx, 5kx, 10kx and 25kx. High magnification (25,000x) calibrated TEM micrographs (5 per MCL) were used for the quantification of collagen fibril diameter using ImageIQ (IQ-bot) automated software by ImageIQ, Inc. Cleveland, OH. The total number of fibril (count) that were analyzed per MCL is shown in ***Table 1***. Mean fibril diameter measurements were combined for basic statistical comparison between treated, untreated and normal control and reported as means ± standard error (SE), followed by a one-way analysis of variance (ANOVA), where *p < 0*.*05* was considered statistically significant.

**Table 1.** Fibril Count and Mean Fibril Diameter (nm) expressed as Mean ± SE

| Rabbit ID | Time Point Post Procedure | Matrix | Total Fibril Count | Mean ± SE |
| --- | --- | --- | --- | --- |
| 603 Normal MCL | 2 weeks | Native – Normal Control | 5,847 | 92 ± 0.3 |
| 603 Left MCL | 2 weeks | Untreated Gap Defect | 13,259 | 50 ± 0.2 |
| 603 Right MCL | 2 weeks | Treated Gap Defect | 9,599 | 75 ± 0.2 |
| 608 Left MCL | 6 weeks | Normal – Uninjured | 7,043 | 98 ± 0.5 |
| 608 Right MCL | 6 weeks | Treated Gap Defect | 4,194 | 88 ± 0.6 |
| 740 Left MCL | 6 weeks | Normal – Uninjured | 7,164 | 92 ± 0.3 |
| 740 Right MCL | 6 weeks | Untreated Gap Defect | 13,844 | 62 ± 0.2 |

## Results

### Collagen fibril distribution and statistical analysis

#### Rabbit 603

At 2 weeks post procedure, TEM analysis of collagen fibril diameter for the **native MCL** (*normal control*), displayed a bimodal distribution of fibrils with mean diameter of 92 ± 0.3 *nm*. The **right MCL** *(treated gap defect*) presented a bimodal fibril distribution consisting of a population of smaller fibrils relative to that observed in normal control with a mean diameter of 75 ± 0.2 *nm*. In contrast, the **left MCL** (*untreated gap defect*), showed a unimodal fibril distribution composed of a homogeneous population of small fibrils with a mean diameter of 50 ± 0.2 *nm*. Importantly, side-to-side comparison showed a statistically significant (*p < 0*.*0001*) difference in the mean fibril diameter in the treated (Right) MCL relative to untreated (Left) MCL (**Figures 1 and 2**).

**Figure 1.**
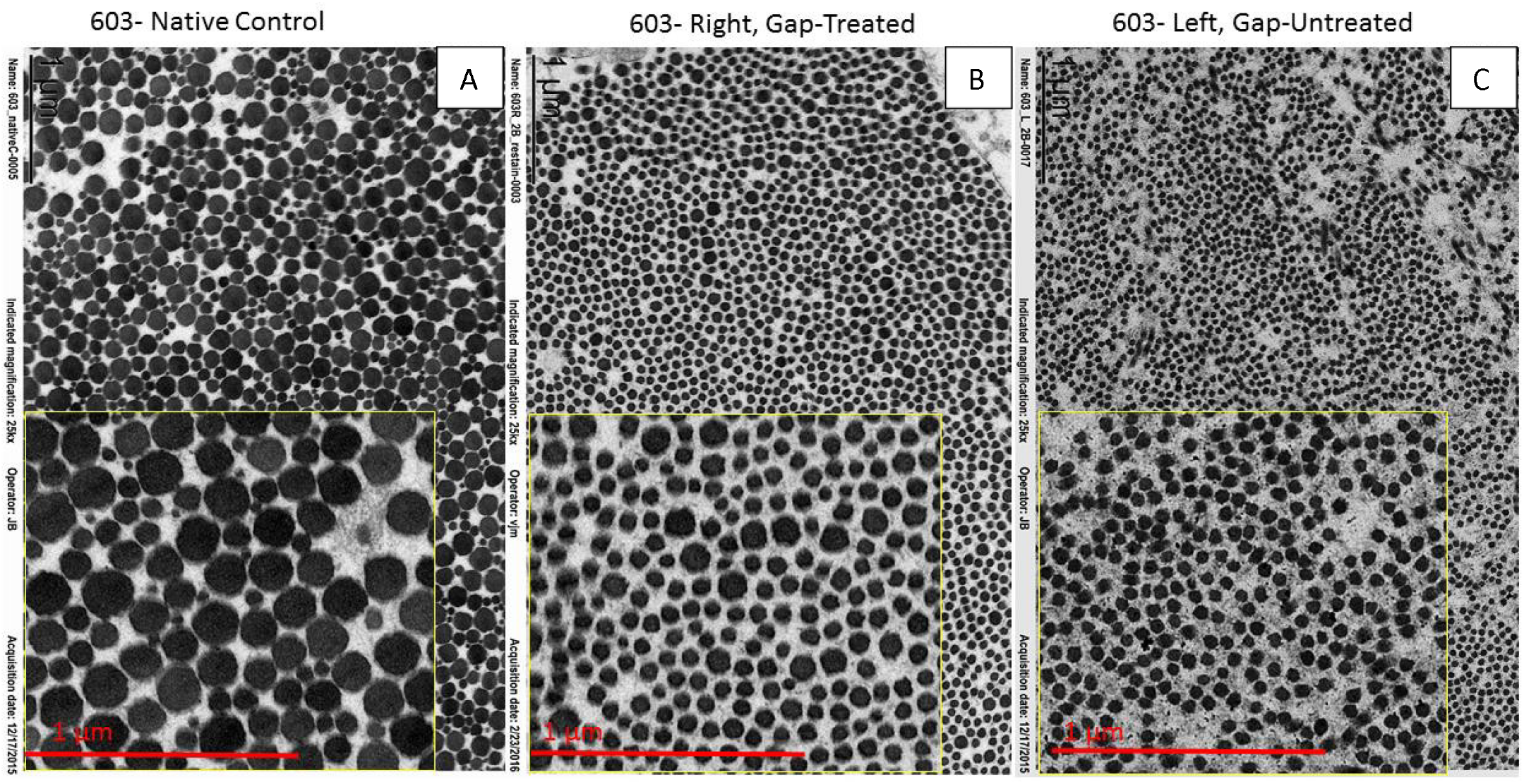
Rabbit 603: Representative high magnification (25kx) TEM micrographs of collagen fibrils cross-sectional view; **(A)** from native (*normal*) control showing a bimodal population of collagen fibrils, typically found in normal, mature MCL. **(B)** Right MCL (*treated gap defect*) showed similar bimodal population of collagen fibrils but significantly smaller than the native control but significantly larger than untreated MCL (p <0.0001). **(C)** Left MCL (*untreated gap defect*) showing unimodal, homogeneous population of small collagen fibrils. Note, inset a higher magnification showing cross-section of collagen fibrils.

**Figure 2.**
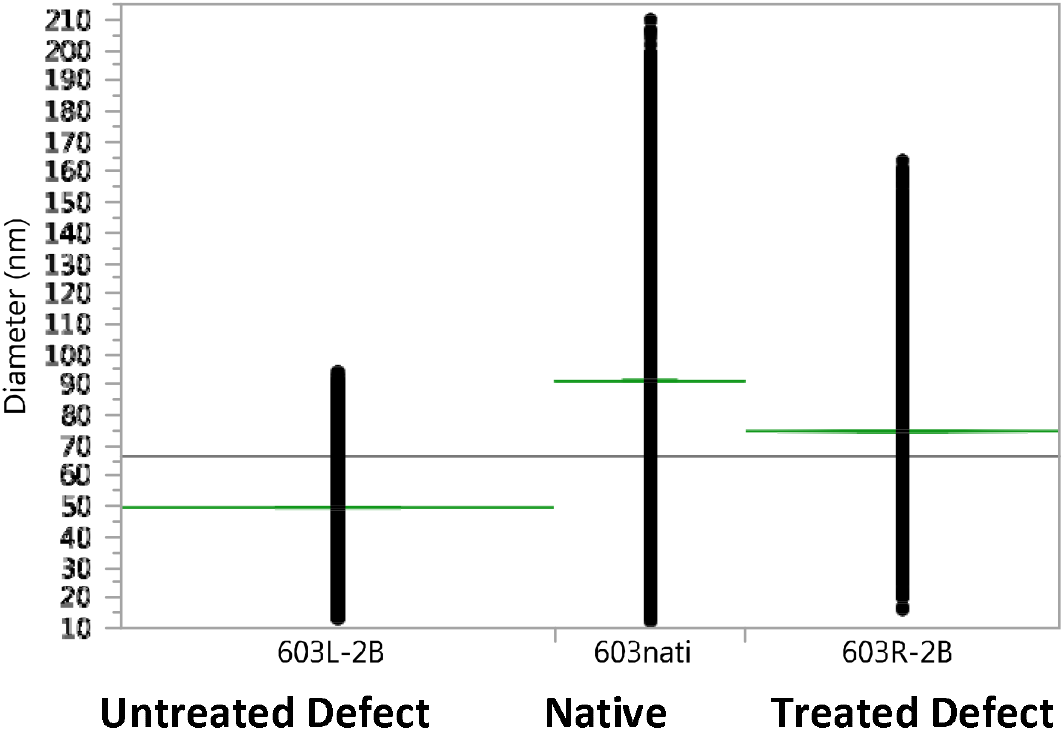
Histogram showing mean fibril diameter of all MCLs studied at 2-weeks post (gap defect) procedure. Side-to-side comparison of MCL gap defect showed statistically significant differences in mean fibril diameter (*p*<0.0001), when comparing treated (75 ± 0.2) vs. untreated (50 ± 0.2) MCL.

#### Rabbit 608

At 6 weeks post procedure, TEM analysis of collagen fibril diameter for the **left MCL** (*normal control)*, showed a bimodal distribution of fibrils with mean fibril diameter of 98 ± 0.5 *nm*. The **right MCL** *(treated gap defect*) presented a bimodal fibril distribution consisting of small and large diameter fibrils with a mean diameter of 88 ± 0.6 *nm*, where *p value < 0*.*0001* for the side-to-side comparison with the left (normal control) MCL of the same animal (Table 1). **Rabbit 740:** At 6 weeks post procedure, collagen fibril diameters for the **right MCL** *(untreated gap defect*) showed a unimodal distribution of a homogeneous population of small fibrils with a mean diameter of 62 ± 0.2 *nm*. The contralateral **left MCL** (*normal control)*, showed a normal bimodal distribution of collagen fibrils with a mean fibril diameter of 92 ± 0.3 *nm*. Inter-animal mean fibril diameter comparison between **right MCL** (*treated gap defect*) in **rabbit 608** (88 ± 0.6 *nm*) and **right MCL** (*untreated gap defect*) in **rabbit 740** (62 ± 0.2 *nm*), demonstrated a statistically significant increase in fibril diameter, where *p<0*.*0001* (**Figures 3, 4 and 5**).

**Figure 3.**
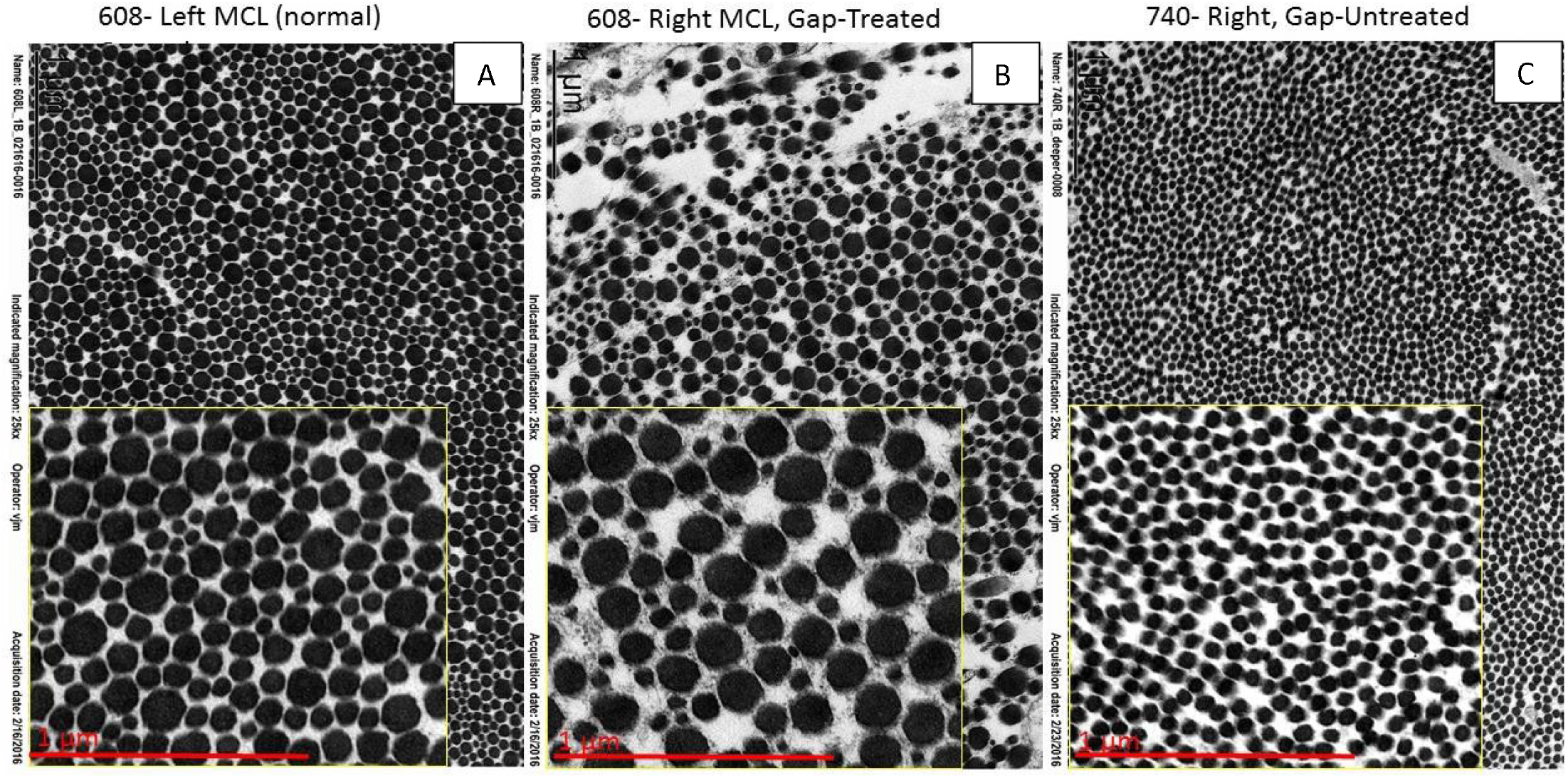
Representative high magnification (25kx) TEM micrographs of collagen fibrils cross-sectional view; Rabbit 608: **Left MCL** native (*normal*) control showing typical bimodal population of collagen fibrils. **(B)** Rabbit 608: **Right MCL** (*treated gap defect*) showing a bimodal population of fibrils with a slight increase of larger fibril diameter and a decrease in fibril distribution/density when compared to (A) normal control. **(C**) Rabbit 740: **Right MCL** (untreated gap defect) showing unimodal distribution of smaller and homogeneous fibrils. Note, inset a higher magnification showing cross-section of collagen fibrils.

**Figure 4.**
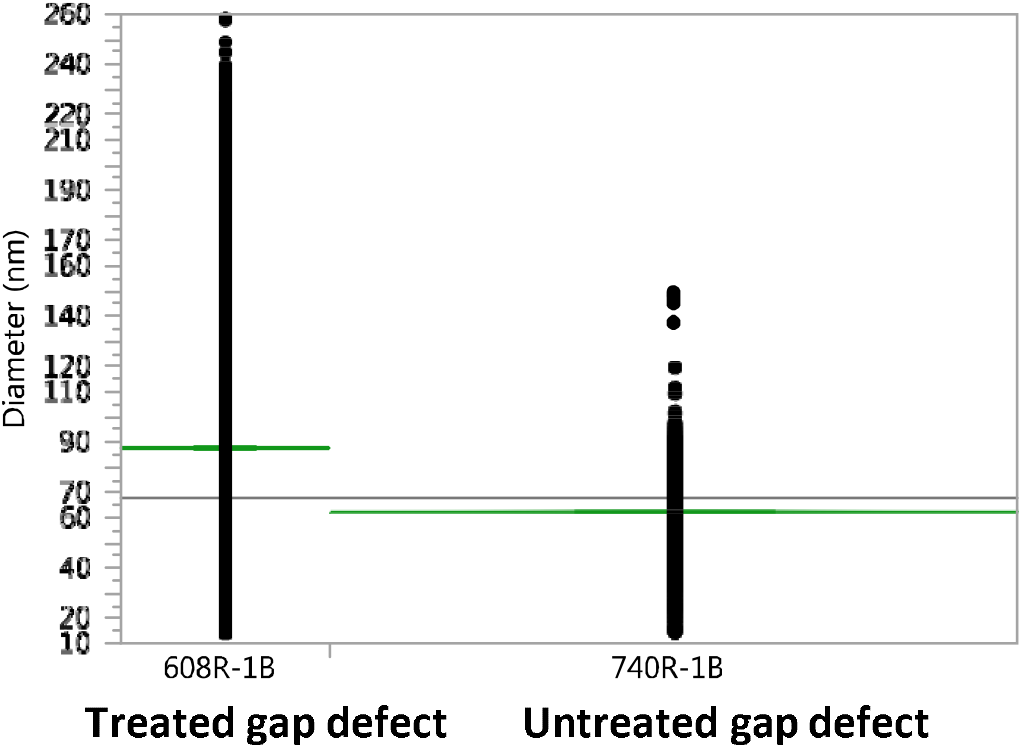
Histogram showing mean fibril diameter of all MCLs studied at 6-weeks post (gap defect) procedure. Inter-animal comparison shows statistically significant differences (*p* <0.0001), when comparing 608 Right MCL (*treated gap defect*) vs. 740 Right MCL (untreated gap defect).

**Figure 5.**
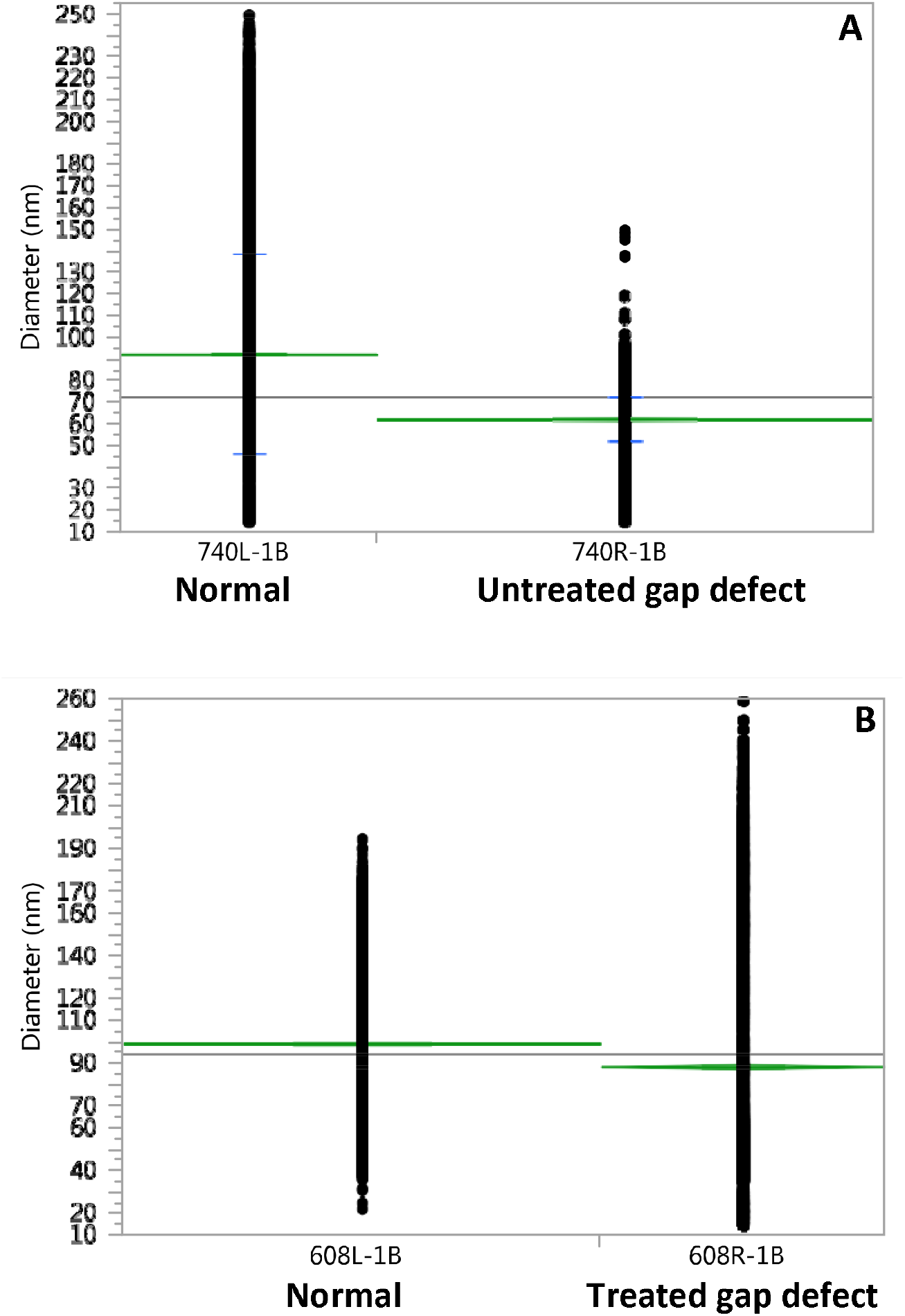
Histogram showing mean fibril diameter of all MCLs studied at 6-weeks post procedure. Results expressed as mean ± SE. (A) Side-to-side comparison of rabbit 740, left normal MCL (92 ± 0.3) vs. right (untreated gap defect) MCL (62 ± 0.2) showed statistically significant differences in mean fibril diameter (p <0.0001). (B) Side-to-side comparison in rabbit 608, left normal MCL (98 ± 0.5) vs. right (treated-gap defect) MCL (88 ± 0.6) showed statistically significant differences in mean fibril diameter (p <0.0001).

### Analysis for Collagen Fibril Distribution

In the 2-week time point (rabbit 603), the native MCL control displayed a broad distribution of fibril diameters with a more heterogeneous population typical of normal uninjured MCL. The untreated left MCL presented a more homogeneous and narrow distribution range of small diameter fibrils. In contrast, the contralateral (treated gap defect) right MCL was characterized by a broader and more heterogeneous distribution of fibril than that observed in the left (untreated gap defect) MCL (**Figure 6**).

**Figure 6.**
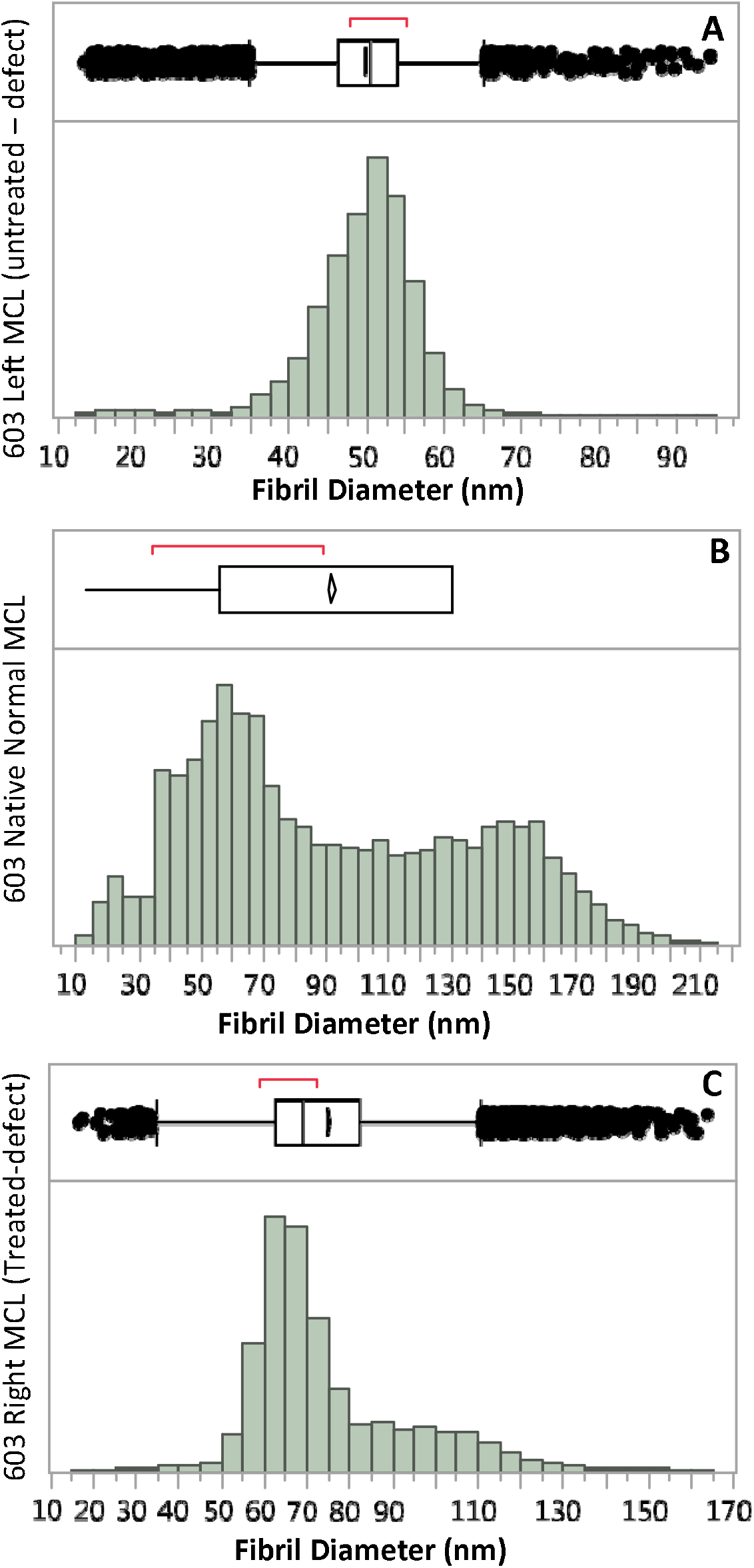
Collagen fibril distribution at 2-weeks post procedure (rabbit 603): (**A**) **left MCL** (untreated gap defect), displaying narrow distribution of small-diameter fibrils consistent with scar repair. (**B**) **native MCL** control displaying a broad distribution of fibril diameter with a more heterogeneous population, typical of normal uninjured MCL. (**C**) **right MCL** displaying a broader distribution of a heterogeneous population of fibril diameters when compared to the left MCL untreated (gap defect).

At 6 weeks post procedure, inter-animal comparison of collagen fibril distribution for normal (uninjured) left MCL from rabbits 608 and 740 showed both animals presented a similar broad range of fibril distributions with a heterogeneous population of fibril diameters. In contrast, in rabbit 608 the right MCL (treated gap defect) presented a broader, more heterogeneous population of fibril diameters, relative to rabbit 740 where the right MCL (untreated gap defect) displayed a narrower distribution characterized by a small homogeneous population of fibrils (**Figure 7**).

**Figure 7.**
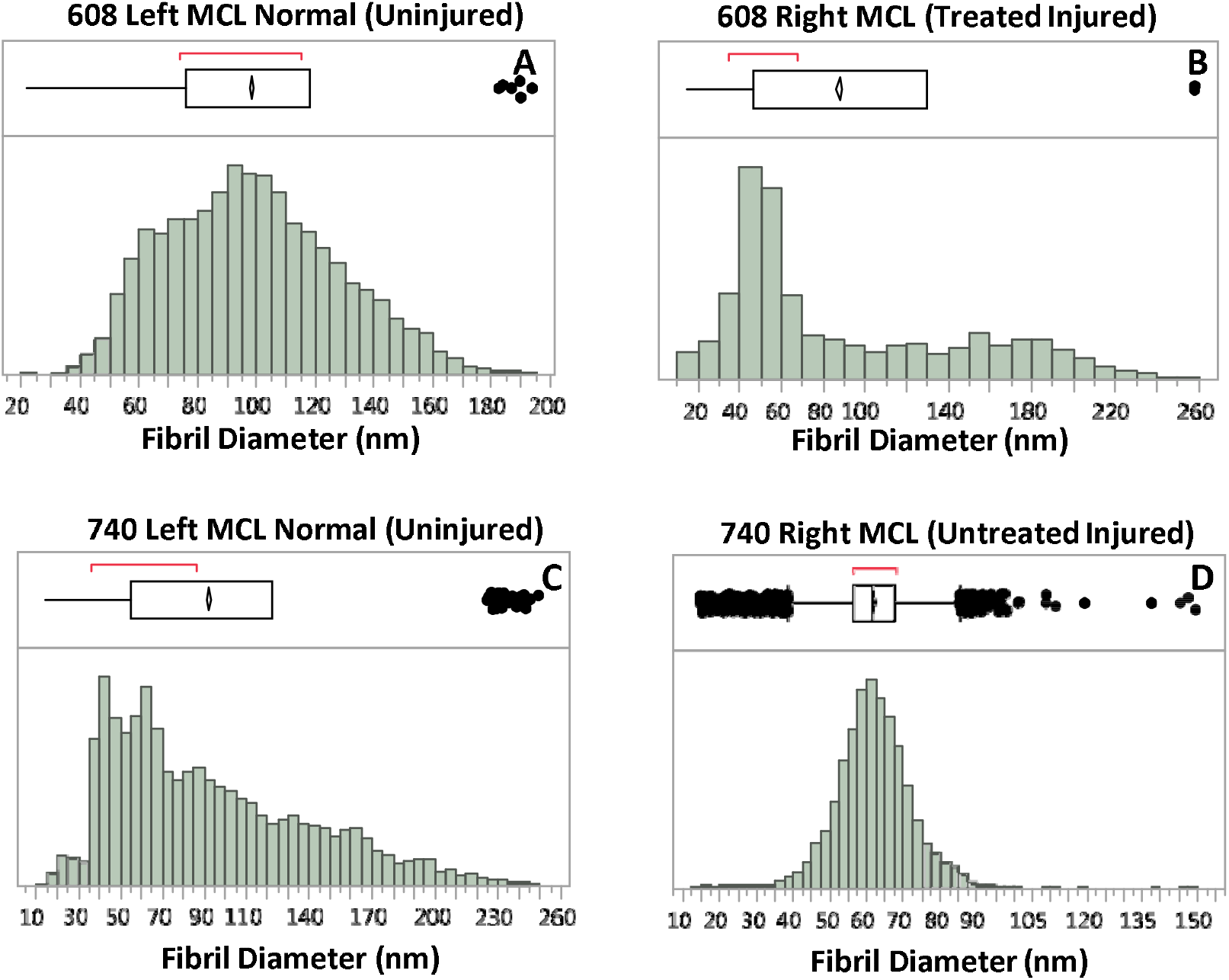
Collagen fibril distribution at 6-weeks post procedure (rabbits 608 and 740): (**A and C)** normal uninjured left MCLs, both showing a broad distribution of fibrils with heterogeneous population in diameter, typical of normal uninjured MCL. (**B and D**) The right MCLs showing a broad distribution of a heterogeneous population of fibril diameters in 608 right MCL (treated gap defect) and a narrow range in fibril distribution with small diameter in 740 right MCL (untreated gap defect).

Collectively, biologically augmented MCL defects demonstrated substantially greater collagen fibril maturation than untreated defects. At 2 weeks, mean collagen fibril diameter was approximately 50% greater in treated versus untreated MCL (75 vs. 50 nm; p<0.0001). At 6 weeks, mean fibril diameter was approximately 42% greater in treated versus untreated MCL (88 vs. 62 nm; p<0.0001). By 6 weeks, treated ligament had recovered to approximately 90% of the mean fibril diameter observed in native, uninjured MCL (88 vs. 98 nm).

## Discussion

In all animals, the normal control MCL (left MCL for rabbits 608 and 740; MCL adjacent to the injury site for rabbit 603) displayed a typical bimodal distribution of small and large diameter fibrils without showing a statistically significant difference in mean fibril diameter when compared between 2 weeks and 6 weeks post procedure. None of the untreated gap defects contained the fibril populations typical of uninjured (normal) rabbit MCLs. These observations are consistent with previously reported TEM studies by Frank et al. [1-3].

More importantly, there was a statistically significant increase in mean fibril diameter between treated and untreated MCL at 2-weeks and 6-weeks post gap defect procedure (*p<0*.*0001*). This is a remarkable finding as these changes can be correlated to histological studies (data not shown), where the regenerated ligament area (gap defect) was shown to be characterized by adequate tissue structure and cellular organization comprised of collagen-rich matrix with longitudinally arranged collagen fibers containing abundant interspersed fibroblasts.

Interestingly, while both of the treated (gap-defect) right MCL (rabbit 603 and 608), displayed a bimodal population of collagen fibrils, there was an increase in mean fibril diameter at 6 weeks (88 ± 0.6 nm) compared to 2 weeks post procedure (75 ± 0.2 nm). These findings suggest that treated MCL are undergoing early healing events that are consistent with tissue regeneration characterized by an increase in fibril diameter and density over time. In contrast, the untreated (gap-defect) MCL showed significantly smaller diameter collagen fibrils (likely type III) with unimodal distribution, which are prevalent during early stages of healing, remodeling and subsequent scar formation. Furthermore, analysis of fibril diameter distribution also indicates that there is a shift in mode from a narrow range of small diameter fibrils in the untreated, injured MCL to a broader range of larger diameter fibrils in the treated MCL regardless of the time point.

These findings suggest a qualitative distinction between regenerative and reparative healing. Untreated ligament defects remained dominated by a relatively homogeneous population of small-diameter collagen fibrils characteristic of scar-associated repair. In contrast, biologically augmented defects developed progressively larger collagen fibrils together with a broader, bimodal fibril distribution that more closely resembled native, uninjured ligament. Thus, treatment was associated not simply with formation of replacement tissue, but with restoration of collagen architecture toward a more native ligament phenotype.

## Conclusion

Using an established rabbit MCL gap-defect model, this study demonstrated a significant regenerative response following biologic augmentation with XWRAP^®^ human amniotic membrane in combination with an amniotic-fluid-derived biologic. Treated ligament defects developed significantly larger collagen fibrils than untreated defects at both 2 and 6 weeks (p<0.0001), with fibril diameter increasing from 75 nm at 2 weeks to 88 nm at 6 weeks. By 6 weeks, treated fibrils reached approximately 90% of the mean diameter observed in native, uninjured ligament. Treated defects also developed a broader, bimodal collagen fibril distribution more closely resembling native MCL architecture, whereas untreated defects remained characterized by a relatively homogeneous population of smaller fibrils associated with reparative/scar healing. Collectively, these findings demonstrate that biologic augmentation with XWRAP^®^ in combination with an amniotic-fluid-derived biologic was associated with progressive restoration of collagen architecture toward a native ligament phenotype.

